# Oscillatory coupling between thalamus, cerebellum and motor cortex in essential tremor

**DOI:** 10.1101/2024.11.11.622917

**Authors:** Alexandra Steina, Sarah Sure, Markus Butz, Jan Vesper, Alfons Schnitzler, Jan Hirschmann

## Abstract

**Background:** Essential tremor is hypothesized to emerge from synchronized oscillatory activity within the cerebello-thalamo-cortical circuit. However, this hypothesis has not yet been tested using local field potentials directly recorded from the thalamus alongside signals from both the cortex and cerebellum, leaving a gap in the understanding of essential tremor.

**Objectives:** To clarify the importance of cerebello-thalamo-cortical oscillatory coupling for essential tremor.

**Methods:** We investigated oscillatory coupling between thalamic local field potentials and simultaneously recorded magnetoencephalography in 19 essential tremor patients with externalized deep brain stimulation electrodes. Brain activity was measured while patients repeatedly adopted a tremor-provoking posture and while pouring rice from one cup to another. In a whole-brain analysis of coherence between the ventral intermediate nucleus of the thalamus and cortex, we contrasted epochs containing tremor and epochs lacking tremor.

**Results:** Both postural and kinetic tremor were associated with an increase of thalamic power and thalamo-cortex coherence at individual tremor frequency in the bilateral cerebellum and primary sensorimotor cortex contralateral to tremor. These areas also exhibited an increase in corticomuscular coherence in the presence of tremor. The coupling of motor cortex to both thalamus and muscle correlated with tremor amplitude during postural tremor.

**Conclusions:** These results demonstrate that essential tremor is indeed associated with increased oscillatory coupling at tremor frequency within a cerebello-thalamo-cortical network, with coupling strength directly reflecting tremor severity.

## Introduction

Upper limb action tremor is the main symptom of essential tremor, the most prevalent movement disorder worldwide.^1^ Deep brain stimulation (DBS) of the ventral intermediate nucleus of the thalamus (VIM) is an effective therapy for severe essential tremor. The insertion of electrodes for DBS provides the unique opportunity to record signals directly from the VIM.

Intraoperative studies have identified tremor-synchronous bursting cells in the VIM^2^ and local field potential (LFP) recordings have uncovered oscillations at tremor frequency, and double tremor frequency, coherent with muscle activity in the tremulous arm.^3^

Apart from the VIM, other parts of the brain, such as cerebellum and motor cortex, have been implicated in the pathophysiology of essential tremor. Studies using functional magnetic resonance imaging (fMRI) have found tremor-related BOLD signal fluctuations in cerebellum, thalamus and motor cortex.^4,5^ Further, non-invasive electroencephalography (EEG) and magnetoencephalography (MEG) studies have revealed tremor-synchronous activity in cerebellum and primary motor cortex.^6–8^

Based on these findings, it is often assumed that essential tremor emerges through synchronized activity within the cerebello-thalamo-cortical circuit, even though tremor-related synchronization of thalamic, cortical and cerebellar oscillations has rarely been investigated so far. Two case studies describe coherence between thalamus and motor cortex, ^9,10^ but a group-level, brain-wide analysis is lacking, as is evidence for tremor-related coupling between thalamus and cerebellum.

Studying these network synchronization processes in humans is challenging. While functional magnetic resonance imaging (fMRI) has provided important evidence for the involvement of the cerebello-thalamo-cortical circuit in tremor,^4,5^ it lacks the temporal resolution required to capture the fast dynamics of tremor. MEG and EEG, on the other hand, have sufficient temporal resolution but have limited sensitivity to deep sources, such as the thalamus.

Here, we overcome these difficulties by means of simultaneous recordings from externalized DBS electrodes, MEG and muscle activity in patients with essential tremor. Using this methodology, we provide, to the best of our knowledge, the first description of the network topology of thalamo-cortical coupling, for both postural and kinetic tremor. In addition, we demonstrate the behavioural relevance of thalamo-cortical coupling by relating it to tremor severity.

## Methods

### Patients and recordings

A total of 19 patients with essential tremor undergoing surgery for DBS participated in the study. The study was approved by the Ethics Committee of the Medical Faculty at Heinrich Heine University Düsseldorf (ET: “2018-217-Zweitvotum”, “2021-1587-andere Forschung erstvotierend”) and the patients provided written informed consent before the measurement, according to the Declaration of Helsinki.

The recordings took place the day after implantation of DBS macroelectrodes, before the pulse generator was implanted. This allowed for the recording of LFPs from externalized leads. Detailed patient information is summarized in (**Table 1)**.

**Table 1.** Patient details.

| **Patient ID** | **age [y]** | **sex** | **disease duration [y]** | **electrode type** |
| --- | --- | --- | --- | --- |
| ET01 | 65 | m | 19 | Abbott Infinity |
| ET02 | 69 | m | 18 | Abbott Infinity |
| ET03 | 71 | m | 20 | Abbott Infinity |
| ET04 | 60 | f | 49 | Abbott Infinity |
| ET05 | 62 | m | 50 | Abbott Infinity |
| ET06 | 65 | m | 30 | Boston Sc. Vercice standard |
| ET07 | 58 | m | 5 | Abbott Infinity |
| ET08 | 77 | f | 8 | Boston Sc. Vercice Cartesia |
| ET09 | 74 | m | 20 | Abbott Infinity |
| ET10 | 30 | m | 25 | Abbott Infinity |
| ET11 | 57 | f | 51 | Abbott Infinity |
| ET12 | 76 | m | n.a. | Abbott Infinity |
| ET13 | 54 | m | 39 | Medtronic SenSight |
| ET14 | 62 | f | 56 | Abbott Infinity |
| ET15 | 65 | m | 20 | Boston Sc. Vercice Cartesia |
| ET16 | 71 | m | 20 | Boston Sc. Vercice Cartesia |
| ET17 | 67 | m | 15 | Abbott Infinity |
| ET18 | 82 | f | 62 | Boston Sc. Vercice Cartesia |
| ET19 | 68 | f | 35 | Boston Sc. Vercice Cartesia |
| $\mu \pm \sigma$ | $65 \pm 11$ | | $31 \pm 20$ | |

y: year, m: male, f: female, n.a.: not available, $\mu$ : mean, $\sigma$ : standard deviation.

LFPs were measured in combination with MEG, EMGs from both forearms (extensor digitorum communis and flexor digitorum communis), accelerometer signals from both index fingers, and electrooculograms. Intracranial LFPs were recorded from bilateral electrodes targeting the VIM and were referenced to a mastoid reference. MEG signals were recorded by a 306-channel MEG system (Vectorview, MEGIN) with a sampling rate of 2 kHz.

### Paradigm

The experiment consisted of three motor tasks, which were performed following a 5 – 10 min resting state recording, analysed in our previous work.^11^ In essential tremor, patients typically experience action tremor with a frequency of 4-8 Hz, which occurs when maintaining a posture against gravity (postural tremor) or during voluntary movement (kinetic tremor).^12^

In the first motor task (HOLD), patients were instructed to place their elbows on a table in front of them and to elevate both forearms with palms facing inward and fingers spread. This task was carried out for 7 min in total. To avoid fatigue, we alternated holding and resting every 20 s. This task was intended to provoke postural tremor.

The second task (POUR) was designed to provoke kinetic tremor. Patients kept one plastic cup in each hand for the entire task, one filled with rice and the other empty. For this task, we positioned a screen in front of the patients, displaying a red fixation cross. They were instructed to start pouring the rice from one cup into the other, standing on the table, once the fixation cross turned green (Go cue), and to keep the pouring posture until the cross turned red again (Stop Cue). At this point, both cups were to be placed on the table until the next Go cue appeared. The Go and Stop cue were displayed for 10 s and 5 s, respectively. This task was performed in blocks of 2.5 min and each patient completed 2-3 blocks. Due to fatigue, this task was performed by only eight out of 19 patients.

### Data preprocessing

Preprocessing and further analysis steps were performed with the FieldTrip toolbox,^13^ MNE-Python^14^, and custom written MATLAB (the MathWorks) scripts.

We scanned the raw data for bad MEG, LFP, and EMG channels and excluded these from further analyses. Next, we applied temporal signal space separation to the MEG data using MNE-Python’s *mne*.*preprocessing*.*maxwell_filter* in order to reduce artefacts. We set *st_duration* to 10 s and *st_correlation* to 0.98.

The rest of the analysis was performed with the FieldTrip toolbox. The data were down-sampled to 200 Hz and only the 204 planar gradiometers were used for further analysis. LFPs were rearranged into a bipolar montage by subtracting the signals of adjacent contacts. EMGs were high-pass filtered at 10 Hz and full-wave rectified.

### Tremor

We inspected EMG and accelerometer signals to identify tremor and tremor-free epochs. An example EMG trace from one patient is depicted in (**Fig. 1A)**. To avoid any tremor-related activity, we labelled epochs as tremor-free only if we found no indication of tremor in either hand, which was mostly the case for the pauses in between movements. In three cases, tremor persisted in the pauses so that we had to extract tremor-free epochs from the resting-state recordings.

**Figure 1.**
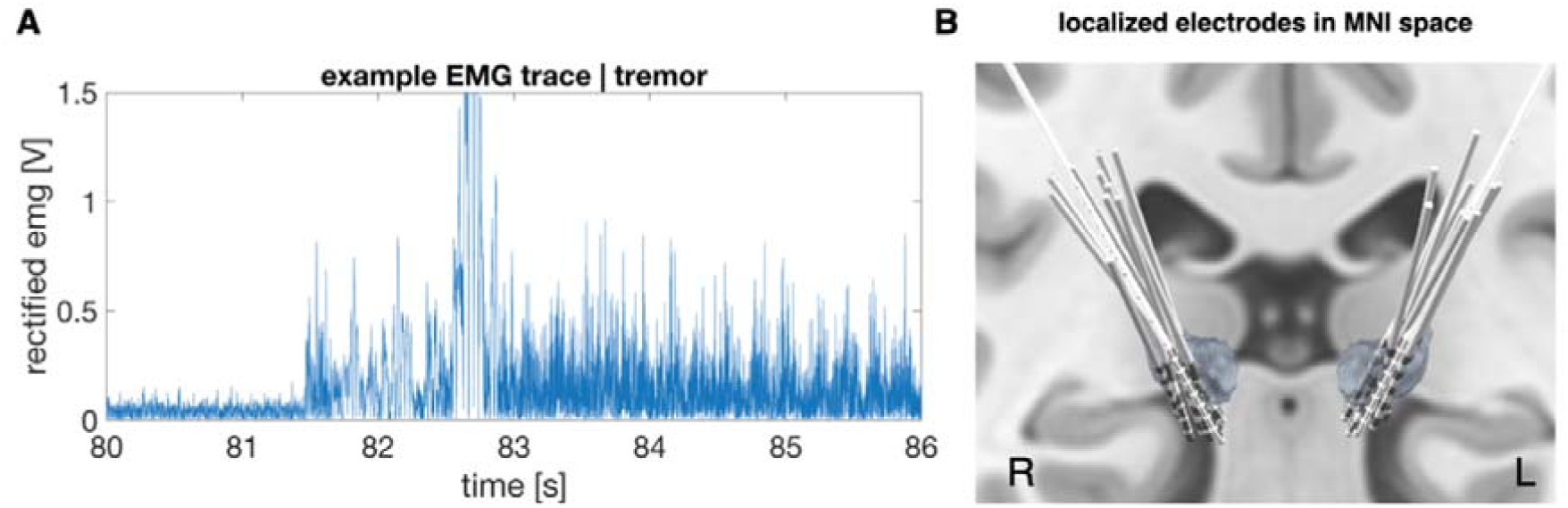
Electromyography signals and deep brain stimulation electrodes targeting the ventral intermediate nucleus of the thalamus. (A) 10 Hz high-pass filtered and rectified and EMG signal during change from rest to hold in one patient. Tremor started immediately after the arm had been lifted. **(B)** Electrodes targeting the VIM, localized with Lead-DBS.

The presence of tremor was evaluated separately for each body side. While this procedure accounts for the independence of left and right upper limb tremor,^15^ it does not stratify the tremor state of the other body side, which may or may not exhibit tremor at the same time. Because the tremor label pertained to one body side only, we limited all tremor analyses to the corresponding (contralateral) hemisphere.

To verify the presence/absence of tremor, we computed the EMG power spectra for each forearm between 1-35 Hz in 0.5 Hz steps and averaged the spectra of flexor and extensor. In these spectra, we identified individual tremor frequency for each body side (see **Table 2** and **Supplementary Table 1**).

**Table 2.**
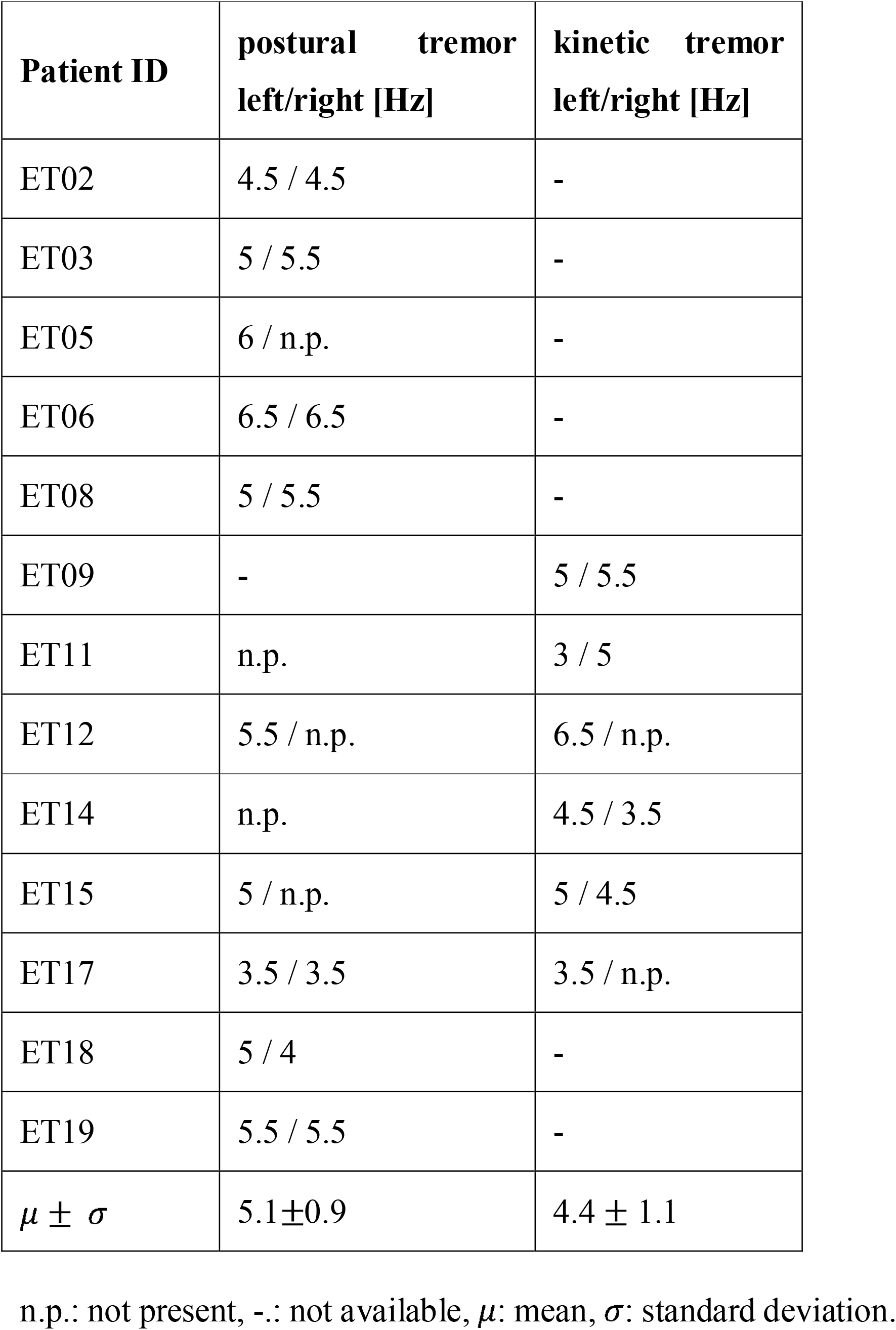
Tremor information only for patients with tremor.

### Power and coherence spectra

For the computation of spectra, we segmented the data into non-overlapping 2 s epochs, convolved the epochs with a Hanning taper, and computed power and thalamo-cortex coherence spectra between 1 and 35 Hz using Welch’s method (frequency resolution: 0.5 Hz).

LFP power spectra are assumed to consist of a periodic component visible as peaks in the spectrum and an aperiodic (1/f) component. We used the fitting oscillations and one over f (FOOOF) algorithm^16^ to model both components and verified a good model fit by visual inspection. For further analysis, only the periodic component was considered.

### Contact localization and contact selection

DBS electrodes were localized with Lead-DBS^17^, using a pre-operative MRI and a post-operative CT scan. The localized electrodes are illustrated in (**Fig. 1B)**. We ensured that electrodes were on target and considered only contacts within the ventral thalamus for further analysis. Additionally, we selected bipolar LFP channels based on their electrophysiological characteristics. For tremor analysis, we selected the channel with the highest power peak at individual tremor frequency contralateral to tremor. Depending on the individual lateralization of tremor, this procedure resulted in either one (one body side affected by tremor) or two selected channels per patient (both body sides affected by tremor). We excluded one patient due to uncertain electrode position.

### Source reconstruction

For each patient, a single-shell head model was generated based on their individual T1-weighted MRI scan (Siemens Mangetom Tim Trio, 3-T MRI scanner, München, Germany). Source reconstruction was performed for a grid with 567 points on the cortical surface, aligned to Montreal Neurological Institute (MNI) space, with a linear constrained minimum variance (LCMV) beamformer.^18^ The regularization parameter *λ* was set to 5%. To account for the rank reduction resulting from temporal signal space separation, we truncated the covariance matrix such that it had the same rank as the Maxwell-filtered data. When computing condition contrasts (tremor *vs*. rest) we applied a common spatial filter to exclude confounds arising due to differences in spatial filters.

#### Source coherence images

We computed thalamo-cortical and corticomuscular coherence spectra in the same way as previously described (see Power and coherence spectra). We averaged activity +/–0.5 Hz around individual tremor frequency and computed one source image per hemisphere in this frequency range.

For epochs containing right hand movement, we mirrored the source images across the midsagittal plane. In consequence, brain activity ipsilateral to movement appears in the left hemisphere, and brain activity contralateral to movement in the right hemisphere in all cases.

### Tremor amplitude

To quantify tremor amplitude, we extracted EMG spectral power at individual tremor frequency +/-0.5 Hz from the 1/f-corrected power spectra and averaged power over flexor and extensor.

## Statistical analysis

As in previous studies,^11,19^ the unit of observation was hemisphere rather than patient (postural tremor: *N*_*hemispheres*_ = 16, kinetic tremor: *N*_*hemispheres*_ = 9). The study had a within-hemisphere design, and we matched the amount of data across conditions for each hemisphere when computing condition contrasts (action tremor *vs*. rest). The statistical analysis was based on nonparametric, two-sample, cluster-based-permutation tests with 1000 random permutations. The tests were two-tailed and the *α* -level was set to 0.05. Multiple comparison correction was implemented by relating all effects to the strongest effects observed in the permuted data (brain-wide or spectrum-wide).^20^ Cortical areas showing differences served as regions of interest for further analyses, such as Pearson correlation between coherence and tremor amplitude.

## Results

### Tremor

In the HOLD task, seven of 19 patients experienced bilateral postural tremor and three patients experienced unilateral tremor. In the POUR task, four of eight patients experienced bilateral kinetic tremor and two patients experienced unilateral tremor (**Supplementary Table 1)**. The average tremor frequency was 5.1 Hz +/– 0.9 Hz (+/–) for postural tremor and 4.4 Hz +/– 1.1 Hz for kinetic tremor.

### Tremor-related thalamic activity

When patients experienced tremor, a clear spectral peak emerged at individual tremor frequency in the electromyography (EMG) power spectrum (cluster-based-permutation-test: postural tremor: *t*_*clustersum*_ = 27.6, *p* = 0.002, (**Fig. 2A**); kinetic tremor: *t* = 19.7, *p* = 0.017, (**Fig. 2B**)), the VIM power spectrum (postural tremor: *t* = 13.0, *p* = 0.009, (**Fig. 2C**); kinetic tremor: *t* = 14.2, *p* = 0.009, (**Fig. 2D**)), and the VIM-EMG coherence spectrum (postural tremor: *t* = 3.1, *p* = 0.15 (not significant), (**Fig. 2E**); kinetic tremor: no cluster, (**Fig. 2F**)), which was absent during rest.

**Figure 2.**
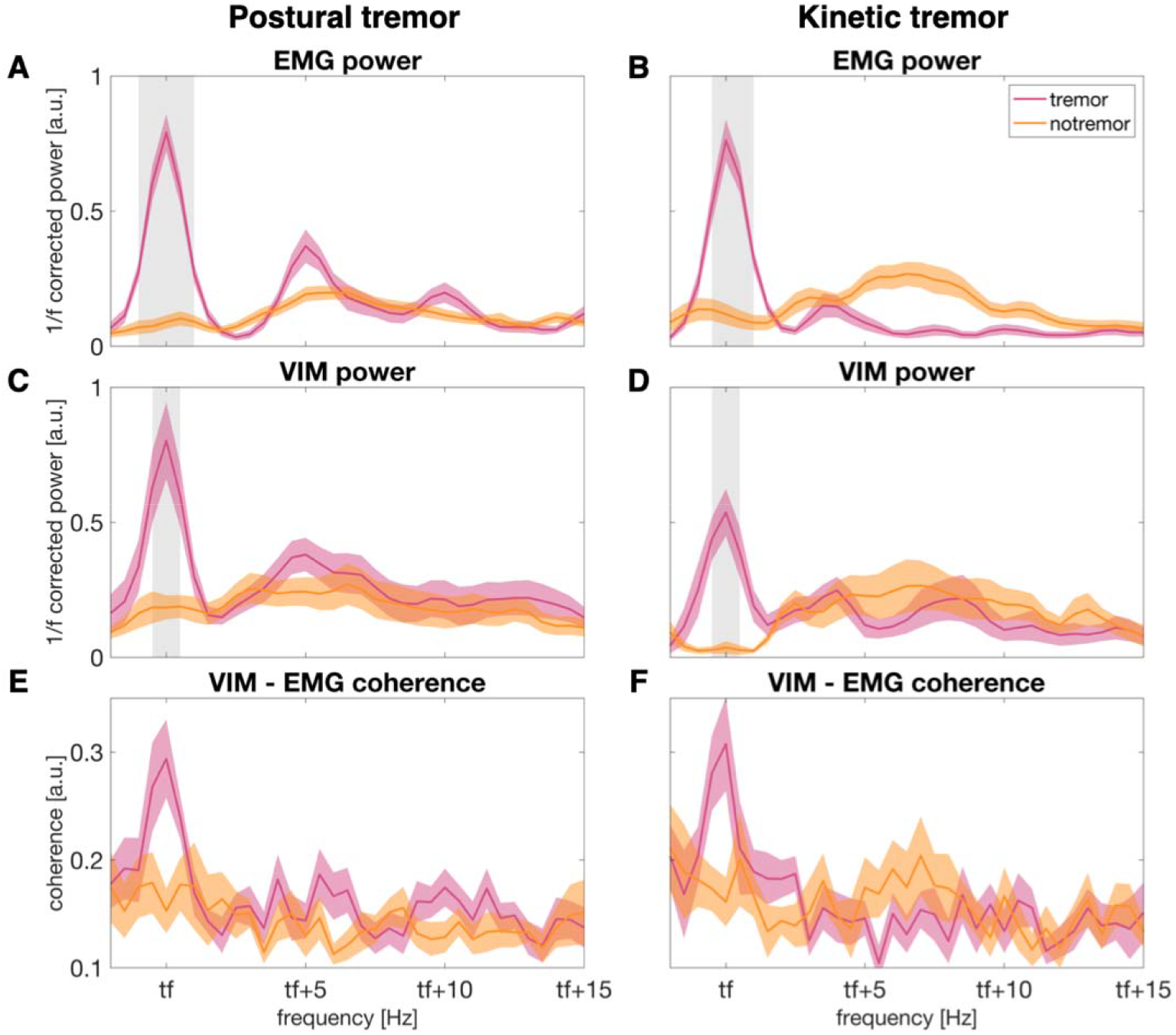
Thalamic and muscle activity during postural and kinetic tremor and tremor-free epochs. Averaged EMG activity of the tremulous arm during postural **(A)** and kinetic **(B)** tremor and tremor-free (no tremor) epochs. 1/f-corrected VIM power contralateral to the tremulous arm during postural **(C)** and kinetic **(D)** tremor. Coherence between the tremulous arm and the contralateral VIM during postural **(E)** and kinetic **(F)** tremor. All spectra were shifted along the frequency axis to align them to individual tremor frequency (tf). For postural tremor, the spectra are averages over 16 hemispheres from nine patients. For kinetic tremor, the spectra are averaged over nine hemispheres from five patients. The shaded areas (pink and yellow) represent the standard error of mean. The grey shadings mark significant differences between tremor and tremor-free epochs. tf = individual tremor frequency.

### Coherence - Postural tremor

Coupling at tremor frequency between cortex and the VIM contralateral to the tremulous arm was stronger in presence than in the absence of tremor. This effect mapped to sensorimotor cortex contralateral to tremor (cluster-based-permutation-test: *t*_*clustersum*_ = 132.23, *p* = 0.002; MNI-coordinates maximal t-value: X = +/-45.4 mm, Y = -30 mm, Z = 62.9 mm), the ipsilateral cerebellum (*t* = 68.19, *p* = 0.008; X = –/+ 8.5 mm, Y = -90 mm, Z = -35 mm) and the contralateral cerebellum (*t* = 39.22, *p* = 0.021; X = +/-25 mm, Y = –50 mm, Z = –60 mm; **Fig. 3A**)).

**Figure 3.**
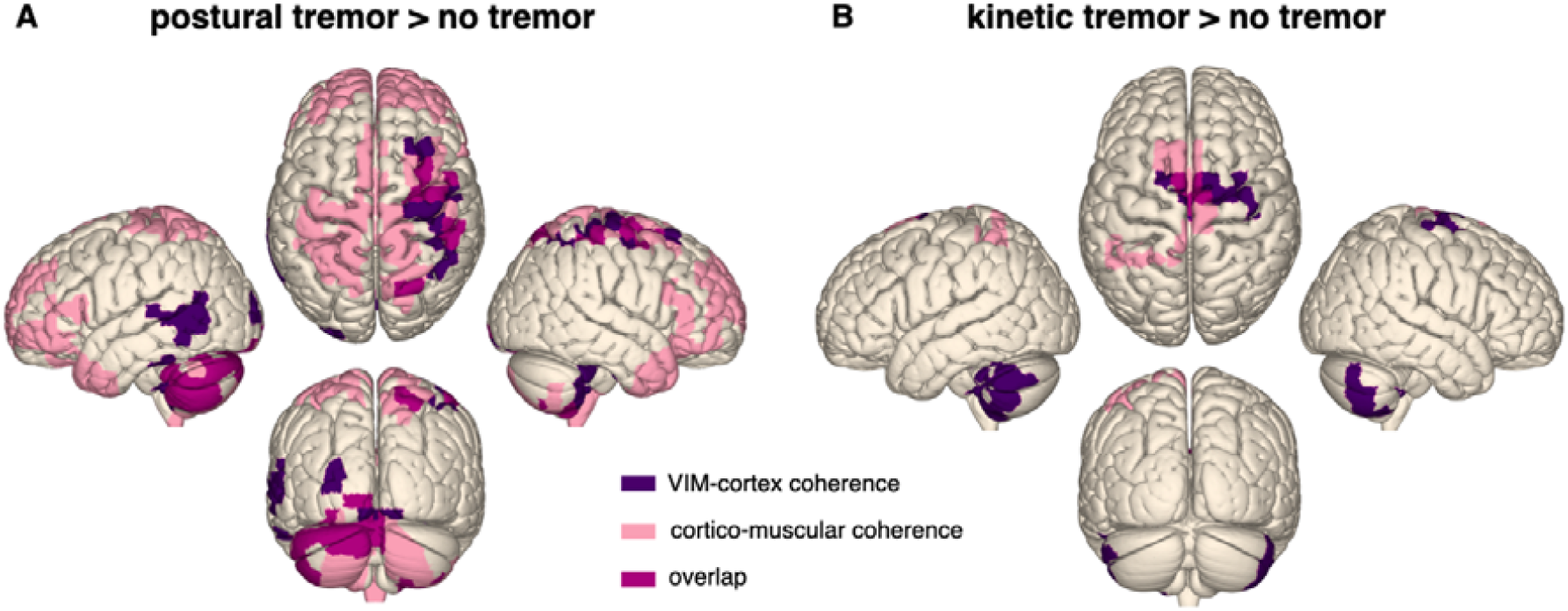
Thalamo-cortical and corticomuscular coherence increased during postural and kinetic tremor. The surface plots illustrate the increase of VIM-cortex (purple) and cortico-muscular coherence (light pink) during postural **(A)** and kinetic **(B)** tremor at individual tremor frequency +/–0.5 Hz. The overlap between VIM-cortex and corticomuscular coherence is displayed in pink. Only coherence to the VIM contralateral to movement is displayed. Left hemisphere: ipsilateral to tremor; right hemisphere: contralateral to tremor.

Cortico-muscular coupling increased in similar regions: bilateral motor cortex (*t* = 205.37, *p* = 0.004; X = +/– 23.7 mm, Y = -60 mm, Z = 70.2 mm), bilateral cerebellum (*t* = 172.83, *p* = 0.006; X = +/–26.6 mm, Y = –90 mm, Z = -31.8 mm), and bilateral prefrontal cortex (*t* = 271.34, *p* = 0.002; X = +/– 40.7 mm, Y = 50 mm, Z = 21.7 mm; **Fig. 3A**)).

VIM-cortex and corticomuscular coherence overlapped in several areas, such as the hand area of sensorimotor cortex contralateral to tremor, as well as in the cerebellum ipsilateral to tremor. Yet, the changes in cortico-muscular coherence were more widespread, including additional frontal and parietal areas.

The VIM-cortex and corticomuscular coherence spectra for the sensorimotor cortex contralateral to tremor, the cerebellum ipsi- and contralateral to tremor are displayed in (**Fig. 4A**) and (**Fig. 5A**).

**Figure 4.**
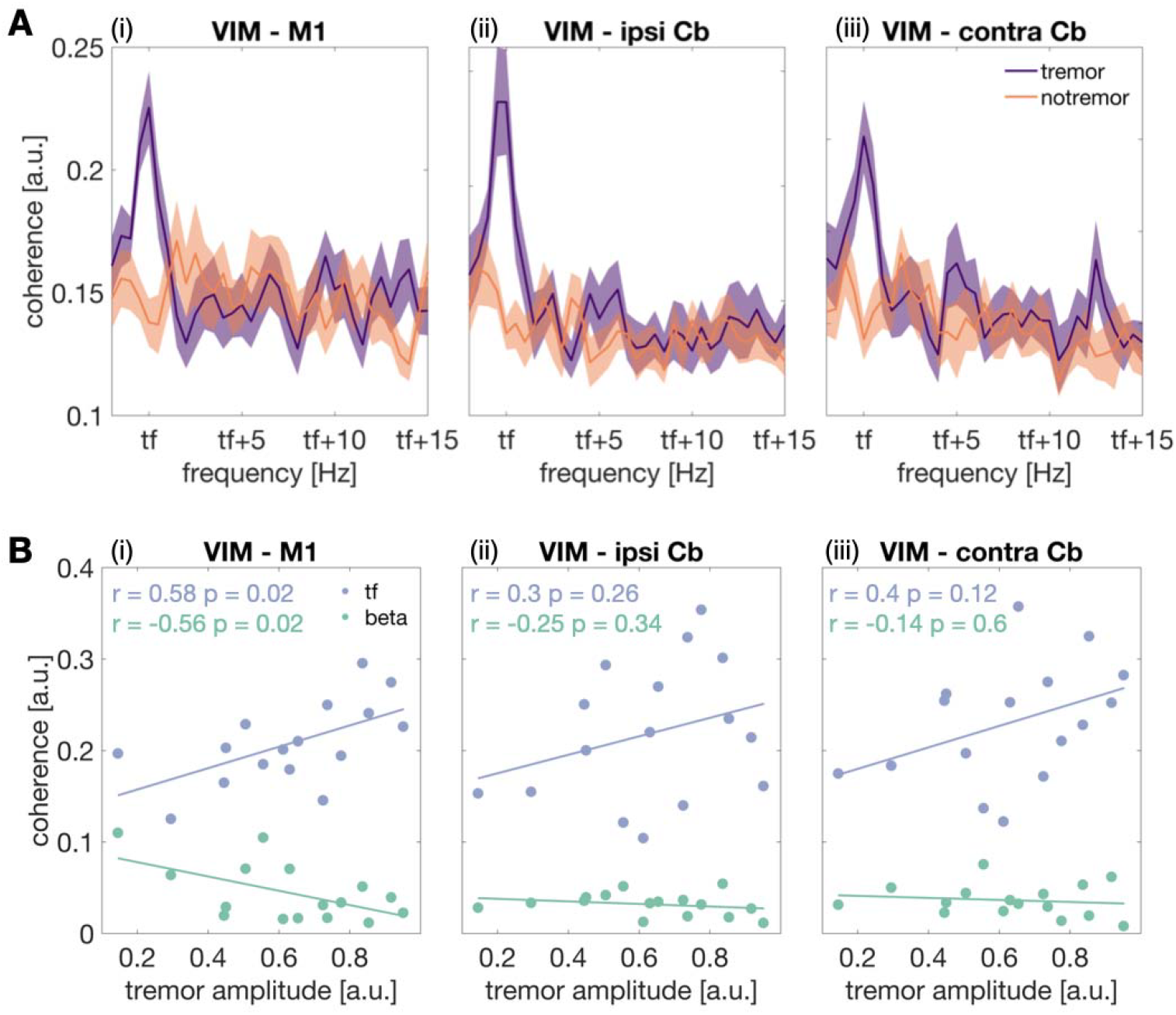
Thalamo-cortex coherence spectra and correlation between postural tremor amplitude and coherence. (A) Coherence between VIM and (i) motor cortex contralateral to tremor, (ii) cerebellum ipsilateral to tremor and (iii) cerebellum contralateral to tremor, for postural tremor and tremor-free epochs (no tremor). Coherence spectra were averaged across patients. Shaded areas represent the standard error of mean. **(B)** Scatter plots illustrate the relationship between tremor amplitude and VIM-cortex coherence at tremor frequency and in the beta band (13-35 Hz) during postural tremor. Cb = cerebellum; M1 = primary motor cortex; ipsi = ipsilateral to tremor; contra = contralateral to tremor; tf = individual tremor frequency.

**Figure 5.**
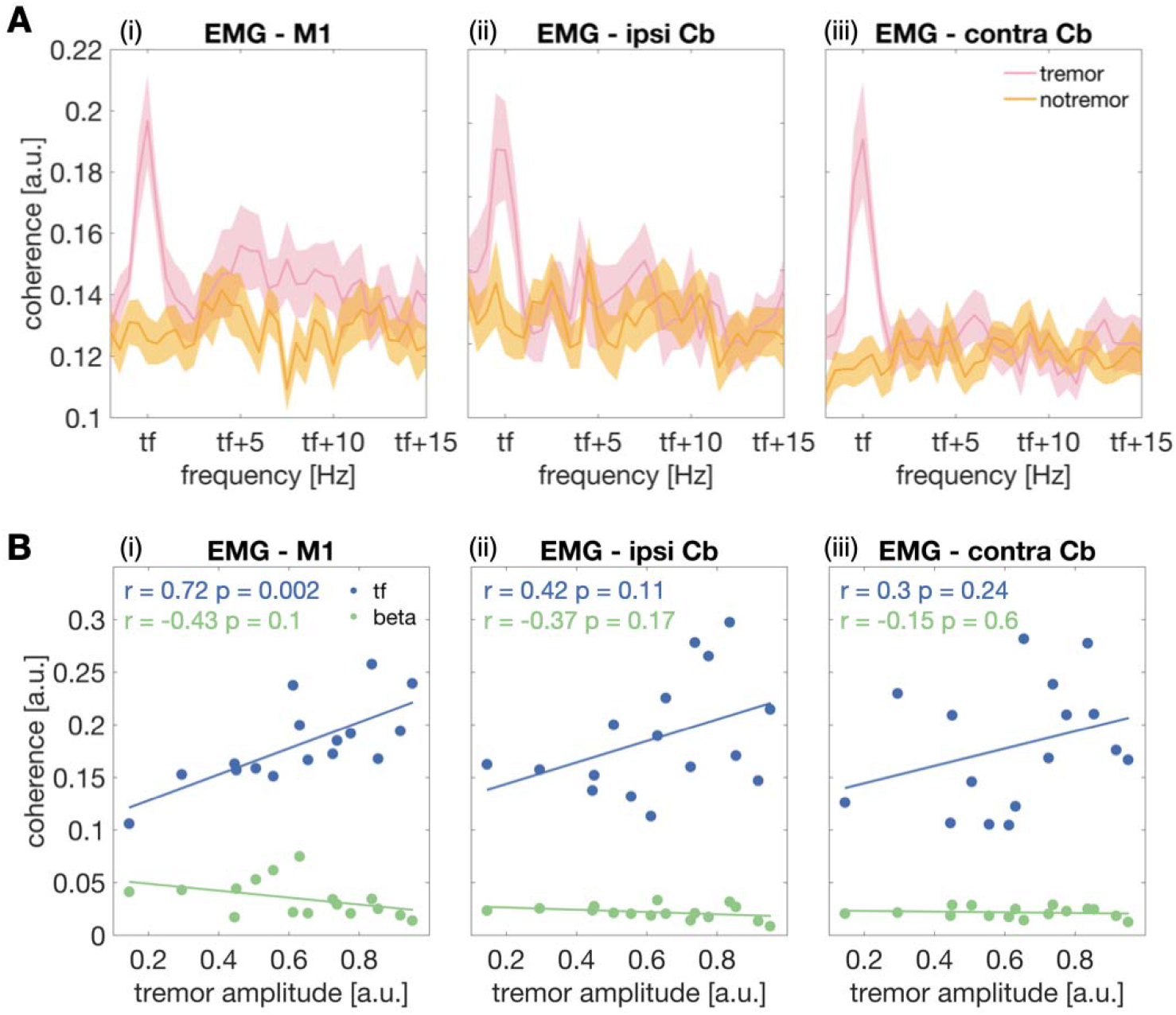
Cortico-muscular coherence spectra and correlation between postural tremor amplitude and coherence. (A) Coherence between EMG and (i) motor cortex contralateral to tremor, (ii) cerebellum ipsilateral and (iii) contralateral to tremor, for postural tremor and tremor-free epochs. Coherence spectra were averaged across patients. Shaded areas represent the standard error of mean. **(B)** Scatter plots illustrate the relationship between tremor amplitude and EMG-cortex coherence at tremor frequency and in the beta band (13 – 35 Hz) during postural tremor. Cb = cerebellum, M1 = primary motor cortex, ipsi = ipsilateral to tremor, contra = contralateral to tremor, tf = individual tremor frequency.

### Coherence – Kinetic tremor

During kinetic tremor, similar changes occurred, but the effects were more circumscribed (**Fig. 3B**). Increases of coherence were observed in supplementary motor cortex contralateral to tremor (*t* = 38.8, *p* = 0.012; X = +/-39 mm, Y = -20 mm, Z = 66 mm), the cerebellum ipsilateral (*t* = 62.6, *p* = 0.002; X = +/-50.6 mm, Y = -40 mm, Z = 41.8 mm), and contralateral to tremor (*t* = 37.84 *p* = 0.012). Cortico-muscular coherence increased in medial sensorimotor regions (*t* = 73.3, *p* = 0.01; X = -/+16.9 mm, Y = 10 mm, Z = 67.1 mm).

### Relationship between VIM-cortex coherence and tremor amplitude – Postural tremor

#### Tremor frequency

The amplitude of tremor correlated with VIM-motor cortex coherence at tremor frequency (*r* = 0.59, *p* = 0.017, (**Fig. 4B (i))**) during postural tremor. The relationship between tremor amplitude and VIM-cerebellar coherence, however, was not significant (cerebellum ipsilateral to tremor: *r* = 0.3, *p* = 0.26, (**Fig. 4B (ii)**), cerebellum contralateral to tremor: *r* = 0.4, *p* = 0.12, (**Fig. 4B (iii))**). Postural tremor amplitude also correlated with EMG-motor cortex coherence (*r* = 0.72, *p* = 0.002, **(Fig. 5B (i))**). Again, the correlation with EMG-cerebellar coherence was not significant (cerebellum ipsilateral to tremor: *r* = 0.42, *p* = 0.1, (**Fig. 5B (ii**)), cerebellum contralateral to tremor: *r* = 0.3, *p* = 0.24, (**Fig. 5B (iii))**).

#### Beta band

We found a negative correlation between tremor amplitude and VIM-motor cortex coherence in the beta band (*r* = -0.56, *p* = 0.025, **Fig. 4B (i)**). The relationship between tremor amplitude and VIM-cerebellar beta coherence, however, was not significant (cerebellum ipsilateral to tremor: *r* = -0.25, *p* = 0.34, **Fig. 4B (ii)**), cerebellum contralateral to tremor: *r* = -0.14, *p* = 0.6, **Fig. 4B (iii)**). The correlation between tremor amplitude and EMG-cortex beta coherence was not significant (motor cortex: *r* = -0.4, *p* = 0.1, **Fig. 5B (i)**), cerebellum ipsilateral to tremor: *r* = -0.37, *p* = 0.17, **Fig. 5B (ii)**), cerebellum contralateral to tremor: *r* = -0.15, *p* = 0.6, **Fig. 5B (iii)**).

## Discussion

In this study, we characterized VIM-cortex coupling during tremor in patients with essential tremor, using intracranial recordings from the VIM, in combination with MEG. During postural and kinetic tremor, VIM power and VIM-cortex coherence increased at individual tremor frequency. This effect was most prominent in primary motor and primary somatosensory cortex ipsilateral to the VIM and the bilateral cerebellum. Cortico-muscular coherence also increased during tremor and exhibited a similar spatial organization. Coupling strength of motor cortex to both VIM and muscle correlated with postural tremor amplitude.

### Localization of tremor-related activity

Using intracranial and MEG recordings, we demonstrate that neuronal oscillations in the ventral thalamus synchronize with motor cortical and cerebellar activity in the presence of tremor. Although this is a common narrative in the tremor literature, no study has, to the best of our knowledge, demonstrated this effect in a larger cohort of ET patients.

Our findings add to a growing body of evidence for a central tremor network underlying essential tremor, gathered through a wide range of techniques, including clinical electrophysiology,^6,7^ fMRI,^4,5^ neuropathology,^21,22^ neurostimulation,^23–26^ and tractography.^27,28^ Studies combining EMG and fMRI have localized tremor-associated brain activity by tracking BOLD signal modulations correlated with slow changes in tremor amplitude.^4,29,30^ Similarly, MEG^7^ and EEG^8^ have been combined with EMG to investigate tremor at a smaller timescale. Across studies, the thalamus, the cerebellum, and primary motor cortex have emerged as major hubs of the essential tremor network. Complementary to these findings, electrophysiology and neuromodulation have uncovered important functional aspects of the cerebello-thalamo-cortical circuit. It has been demonstrated, for example, that phase-locked invasive and non-invasive neurostimulation^23–25^ can intensify or weaken tremor, depending on the phase difference between tremor and stimulation. Such tremor entrainment can be achieved by VIM DBS,^24^ and by non-invasive stimulation of the cerebellum^25^ or motor cortex.^23^ These findings emphasize the importance of rhythmic neural activity synchronized across a distributed tremor network, similar to the findings made in Parkinson’s disease.^31^ A correlation between oscillatory coupling and essential tremor severity, however, has not been demonstrated so far. This is one important contribution of the current study, emphasizing the clinical relevance of thalamo-cortical coupling at tremor frequency.

### Cerebellum

The cerebellum is thought to play a major role in the pathophysiology of essential tremor.^32,33^ In line with this notion, we found that both VIM and muscle activity in the tremulous arm were coherent with the bilateral cerebellum during postural and kinetic tremor. In contrast, previous MEG/EEG studies have found tremor-associated neural activity to be limited to the cerebellar hemisphere ipsilateral to the tremulous arm.^6,7^ One explanation for this difference might be that some of our patients experienced bilateral postural tremor, resulting in bilateral activation of the cerebellum. Notwithstanding bilateral cerebellar activation was also visible during unilateral kinetic tremor (POUR). An involvement of the bilateral cerebellum is plausible based on the structural connections of the VIM: it receives inputs from the contralateral cerebellum via decussating fibres, and, to a minor extend, from the ipsilateral cerebellum via non-decussating fibres of the dentato-rubro-thalamic tract.^34^ Moreover, several studies combining EMG and fMRI reported bilateral cerebellar involvement during unilateral tremor in patients with essential tremor, while only the cerebellum ipsilateral to movement was active during mimicked tremor in healthy individuals.^30,35^ This indicates that the recruitment of both cerebellar hemispheres might be a pathological feature.

### Primary sensorimotor cortex

It is well-established that the primary sensorimotor cortex plays an important role in many types of involuntary movement, such as Parkinsonian tremor^31^ or focal dystonia.^36^ The role of thalamo-sensorimotor cortex coupling in essential tremor, however, is less clear. So far, simultaneous LFP-EEG recordings have been conducted in a total of three patients in two studies. In all three cases, peaks at tremor frequency were observed in the VIM-motor cortex coherence spectra.^9,10^ For coupling between muscle and motor cortex, ambiguous results have been reported. Some studies found increased coupling during tremor,^6,7^ while others found coupling in only a few patients,^37^ and one study reported no coupling at all.^38^ Trying to reconcile these findings, it has been speculated that the involvement of the sensorimotor cortex is intermittent.^39^ In this study, we provide evidence for motor cortical involvement in essential tremor: VIM/EMG-motor cortex coupling increased during both postural and kinetic tremor.

During postural tremor, the strength of VIM-/EMG-motor cortex coupling at tremor frequency, but not VIM-/EMG-cerebellar coupling was correlated with tremor amplitude, underpinning the important role of motor cortex. This notion is supported by previous studies demonstrating that non-invasive stimulation of motor cortex reduces essential tremor amplitude.^40^ Interestingly, similar observations have been made for re-emergent tremor in Parkinson’s disease: transcranial magnetic stimulation of the primary motor cortex, but not the cerebellum, modulated tremor amplitude.^41,42^ In addition, connectivity and network mapping studies have unveiled that VIM-DBS at sites stronger connected to primary sensorimotor cortex was associated with superior tremor improvement.^28,43^ Notably, it has further been reported that sensorimotor cortex leads muscle activity during tremor,^44^ suggesting that the increased synchronization with primary sensorimotor cortex might reflect an active involvement of motor cortex rather than field spread from primary somatosensory cortex.

Additionally, we found that coherence between the VIM and motor cortex in the beta band was inversely correlated with tremor amplitude. A negative association between beta activity and tremor has often been reported for resting tremor in Parkinson’s disease.^31,45^ In the case of essential tremor, previous studies have shown a similar negative correlation between beta activity and tremor within the VIM, ^46^ and our findings extend this relationship to thalamo-cortical coupling. Voluntary movements are likewise associated with a reduction of beta activity and together, these results indicate that tremor and voluntary movements might have common underlying mechanisms.^47^

### Postural vs. kinetic tremor

Action tremor can be divided into different types of tremor such as postural and kinetic tremor and these sub-types can co-occur in a single patient. It remains unclear if the subtypes arise from distinct brain regions.^48^ Combined EMG-fMRI studies found activation of cerebellum, motor thalamus, and motor cortex in different kinds of action tremor, suggesting that the cerebello-thalamo-cortical circuit is involved in the generation of different types of tremor.^29,30^ Our findings indicate that similar networks are involved in both postural and kinetic tremor. However, the cortical distribution of coherence with thalamic activity were more wide-spread for postural tremor than for kinetic tremor. This may be due to postural tremor occurring simultaneously in both body sides in some patients, whereas kinetic tremor was unilateral.

## Limitations

Statistical power might have been limited by the sample size of this study. Due to the postoperative microthalamotomy (stun) effect, uni- or bilateral tremor was present in 12 out of 19 patients during the HOLD task, the POUR task, or both. While this sample size is small in absolute numbers, it is substantially larger compared to previous studies measuring thalamo-cortical coupling in humans (N ≤ 3).^9,10^

Further, from a methodological perspective, it would be desirable to match the motor tasks perfectly, e.g. HOLD with tremor vs. HOLD without tremor or mimicked action tremor *vs*. true action tremor. This was not possible in our cohort because the instruction to keep a static posture or to mimic action tremor inevitably elicits actual tremor.

Another limitation of the study is that we did not distinguish between subgroups of essential tremor, such as “normal” essential tremor and essential tremor plus.^12^ Such a distinction would require a larger and more diverse sample.

## Conclusions

Recording thalamic and cortical activity simultaneously, we have demonstrated that tremor episodes in patients are characterized by synchronized oscillations in the ventral intermediate nucleus, cerebellum and sensorimotor cortex, underpinning the role of the cerebello-thalamo-cortical circuit for the pathophysiology of essential tremor.

## Acknowledgements

ASc and JH are supported by Brunhilde Moll Stiftung. ASc also acknowledges support by the Deutsche Forschungsgemeinschaft (DFG, German Research Foundation) - Project ID 4247788381 - TRR 295.

## Author Contributions

A.S.: Data acquisition, data analysis, visualization, methodology, writing – original draft; S. S.: Data acquisition, conceptualization, writing – review and editing; M.B.: Data acquisition, writing – review and editing; J.V.: Resources, writing – review and editing; A. Sc.: Conceptualization, resources, funding acquisition, writing – review and editing; J.H.: Data acquisition, supervision, methodology, project administration, writing – review and editing.

