## Supplementary for "Oscillatory coupling between thalamus, cerebellum and motor cortex in essential tremor"

**Supplementary material**

Alexandra Steina^1^, Sarah Sure^1^, Markus Butz^1^,
Jan Vesper^2^, Alfons Schnitzler^1^, Jan Hirschmann^1^

**Author affiliations:**

1 Institute of Clinical Neuroscience and Medical Psychology, Medical Faculty, Heinrich Heine University, 40225, Düsseldorf, Germany

2 Department of Functional Neurosurgery and Stereotaxy, Neurosurgical Clinic, Medical Faculty, Heinrich Heine University, 40225, Düsseldorf, Germany

**Supplementary Table 1 Task & tremor information.**

| **Patient ID** | **Tasks performed** | **Postural tremor frequency L/R [Hz]** | **Kinetic tremor frequency L/R [Hz]** | **Postural tremor data length L/R [s]** | **Kinetic tremor data length L/R [s]** |
| --- | --- | --- | --- | --- | --- |
| ET01 | H, P | - | - | - | - |
| ET02 | H | 4.5 / 4.5 | - | 88 / 78 | - |
| ET03 | H | 5 / 5.5 | - | 99 / 40 | - |
| ET04 | H | - | - | - | - |
| ET05 | H | 6 / - | - | 153 / - | - |
| ET06 | H | 6.5 / 6.5 | - | 81 / 80 | - |
| ET07 | H | - | - | - | - |
| ET08 | H | 5 / 5.5 | - | 297 / 213 | - |
| ET09 | P | - | 5 / 5.5 | - | 81 / 72 |
| ET10 | H | - | - | - | - |
| ET11 | H, P | - | 3 / 5 | - | 63 / 109 |
| ET12^a^ | H, P | 5.5 / - | 6.5 / - | 236 / - | 102 / - |
| ET13 | H, P | - | - | - | - |
| ET14 | H, P | - | 4.5 / 3.5 | - | 130 / 146 |
| ET15 | H, P | 5 / - | 5 / 4.5 | 174 / - | 128 / 109 |
| ET16 | H, P | - | - | - | - |
| ET17 | H | 3.5 / 3.5 | 3.5 / - | 102 / 90 | 49 / - |
| ET18 | H | 5 / 4 | - | 69 / 65 | - |
| ET19 | H | 5.5 / 5.5 | - | 87 / 137 | - |

H: Hold, B: Button press, P: Pour.

^a^Left VIM excluded due to uncertain electrode position.
